# Single-cell analysis of human airway epithelium identifies cell type-specific responses to *Aspergillus* and *Coccidioides*

**DOI:** 10.1101/2024.09.09.612147

**Authors:** Alfred T. Harding, Arianne J. Crossen, Jennifer L. Reedy, Kyle J. Basham, Olivia W. Hepworth, Yanting Zhang, Viral S. Shah, Hannah Brown Harding, Manalee V. Surve, Patricia Simaku, Geneva N. Kwaku, Kristine Nolling Jensen, Yohana Otto, Rebecca A. Ward, George R. Thompson, Bruce S. Klein, Jayaraj Rajagopal, Pritha Sen, Adam L. Haber, Jatin M. Vyas

## Abstract

Respiratory fungal infections pose a significant threat to human health. Animal models do not fully recapitulate human disease, necessitating advanced models to study human-fungal pathogen interactions. In this study, we utilized primary human airway epithelial cells (hAECs) to recapitulate the lung environment *in vitro* and investigate cellular responses to two diverse, clinically significant fungal pathogens, *Aspergillus fumigatus* and *Coccidioides posadasii*. To understand the mechanisms of early pathogenesis for both fungi, we performed single-cell RNA sequencing of infected hAECs. Analysis revealed that both fungi induced cellular stress and cytokine production. However, the cell subtypes affected and specific pathways differed between fungi, with *A. fumigatus* and *C. posadasii* triggering protein-folding-related stress in ciliated cells and hypoxia responses in secretory cells, respectively. This study represents one of the first reports of single-cell transcriptional analysis of hAECs infected with either *A. fumigatus* or *C. posadasii*, providing a vital dataset to dissect the mechanism of disease and potentially identify targetable pathways.

**Importance:** Fungal infections in the lungs are dreaded complications for those with compromised immune systems and have limited treatment strategies available. These options are restricted further by the increased prevalence of treatment-resistant fungi. Many studies focus on how our immune systems respond to these pathogens, yet airway epithelial cells remain an understudied component of fungal infections in the lungs. Here, the authors provide a transcriptional analysis of primary human airway epithelial cells stimulated by two distinct fungal pathogens, *Aspergillus fumigatus* and *Coccidioides posadasii*. These data will enable further mechanistic studies of the contribution of the airway epithelium to initial host responses and represent a powerful new resource for investigators.

## Introduction

Pulmonary fungal infections are dreaded, particularly in immunocompromised individuals, due to high mortality rates and limitations in treatment strategies (1, 2). We investigated two clinically relevant fungal pathogens, *Aspergillus fumigatus* and *Coccidioides posadasii*, as they have different cell wall compositions, life cycles, and pathogenic strategies. Importantly, the two pathogens are clinically distinct. *A. fumigatus* is ubiquitous in the environment making immunocompromised patients (*e.g.,* lung transplant or allogenic bone marrow transplant recipients) vulnerable to invasive aspergillosis. The mortality from *A. fumigatus*-related pulmonary disease remains unacceptably high (58%), and rates of *Aspergillus* multidrug resistance continue to increase (3–5). Inhaled conidia germinate into hyphae, a strong virulence trait. This change in morphotype dramatically changes the cell wall composition, exposing more antigenic carbohydrates like β-1,3 glucan and galactomannan when the rodlet and melanin layers are shed. Insights into the mechanisms that lung epithelial cells employ to instruct direct immune cells against this dangerous fungus are thus clinically relevant and will guide new therapeutic strategies.

The dimorphic fungi *Coccidioides immitis* or *C. posadasii* causes coccidioidomycosis and are endemic to the southwestern US, Mexico, and South America, with the historical endemic boundaries expanding with climate change (6, 7), increasing the number of humans and animals at risk of exposure (8–10). Unlike *Aspergillus*, *C. posadasii* produces infection in both immunocompetent and immunocompromised individuals, implying that *C. posadasii* can thwart host defense mechanisms that *A. fumigatus* cannot and that distinct mechanisms are involved in controlling these infections (11, 12). Only 40% of *Coccidioides* infections are symptomatic, and these patients often present with acute or progressive self-limiting pneumonia, including the formation of pulmonary nodules (localized disease). *Coccidioides* is inhaled as arthroconidia, which convert into spherules that release endospores. *Coccidioides* differs from *Aspergillus* in virulence and dose required to cause infection. *Coccidioides* causes disease with inhalation of 1-50 arthroconidia in immunocompetent mice (13), whereas *Aspergillus* requires ∼10^4^-10^5^ CFU to cause disease in immunocompromised (but not immunocompetent) patients and mice, again implying differences in host recognition and responses to these distinct fungal pathogens (14). We hypothesize that these differences provoke distinct and cell type-specific epithelial transcriptional signatures from human lung epithelium that direct the ensuing immune response.

To elucidate the cellular responses and pathogen-specific mechanisms of these pulmonary fungal infections, we utilized human airway epithelial cells (hAECs) as an *in vitro* model to recapitulate the lung epithelium. The hAECs form a pseudostratified epithelial layer, closely mimicking the cellular environment of the human airway (15). In this study, hAECs were cultured and infected with either *A. fumigatus* or *C. posadasii*. Following infection, we isolated the cells and performed single-cell RNA sequencing (scRNA-seq) to obtain a high-resolution view of the transcriptional responses of specific cell types comprising the lung epithelium (16).

This approach enabled us to unveil both common and unique cell-specific responses and stress pathways activated by *A. fumigatus* and *C. posadasii* infection. Our findings revealed that *A. fumigatus* infection primarily affected ciliated cells, inducing protein-folding-related stress, whereas *C. posadasii* infection triggered a hypoxia response in secretory cells, coupled with increased cytokine expression. The distinct cellular responses to *A. fumigatus* and *C. posadasii* infections provide a valuable dataset for understanding the mechanisms of pulmonary fungal diseases and identifying potential therapeutic targets.

## Results

### Experimental Design and hAEC Differentiation and Characterization

To better understand the mechanisms by which fungal pathogens infect and cause disease in the lung, we designed a study wherein we generated *ex vivo* human lung epithelia from basal cells isolated from the same donor, infected differentiated hAECs with either *A. fumigatus* or *C. posadasii*, and then performed scRNA-seq (**Fig. 1A**). To create a robust *in vitro* model of the human airway epithelium, we began by isolating basal cells from human donor lung tissues. These primary basal cells were cultured according to previously published protocols (17, 18). Briefly, we expanded cells in small-airway epithelial cell medium (SAGM) supplemented with various growth factors and inhibitors to promote their expansion (details are provided in the methods section). Basal cells were seeded onto Transwell inserts to establish an air-liquid interface (ALI) culture, which were maintained for approximately 23 days. During this period, the basal cells differentiate into a pseudostratified epithelium that mimics the cellular composition and architecture of the human airway (19, 20).

**Figure 1.**
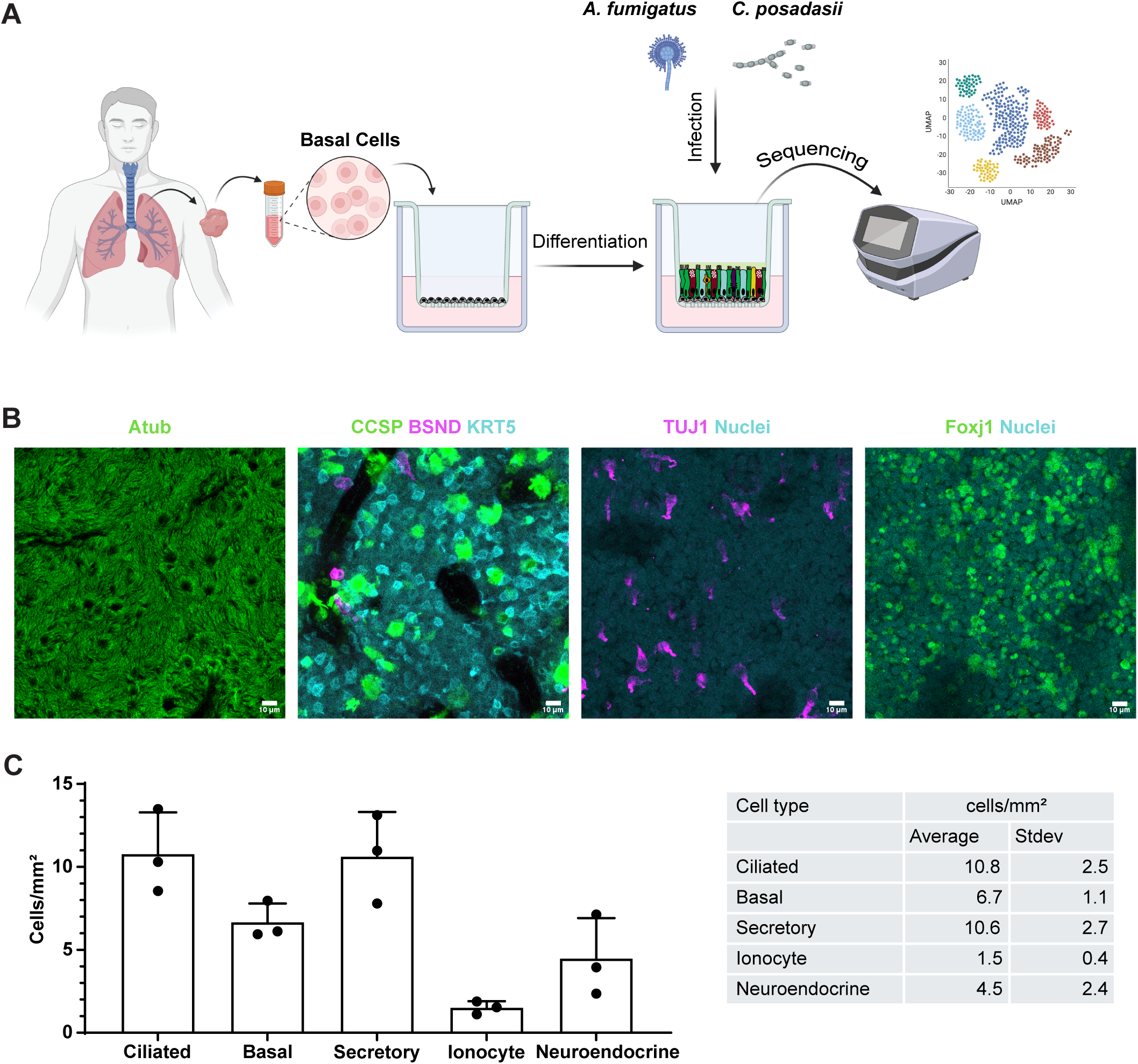
Experimental design and generation of human airway epithelial cell models. **(A)** A schematic outlining the experimental design wherein airway basal cells are isolated from human volunteers, expanded and differentiated, and then infected with either *A. fumigatus* or *C. posadasii* before undergoing scRNA-seq. **(B)** Representative images of fully differentiated hAECs. Cell markers for ciliated cells (AcTub), basal stem cells (KRT5), secretory cells (CCSP), ionocytes (BSND), neuroendocrine (TUJ1), were used. Size bar = 20 μm. **(C)** Quantification of epithelial subtypes per mm^2^ from images captured in (B).

To confirm that our differentiation was successful, we fixed and stained hAECs in a subset of Transwells grown in parallel to ensure the major airway cell subtypes were present. Specific antibodies were used to stain for basal cells (KRT5) (21), club cells (SCGB1A1) (22), and ciliated cells (acetylated tubulin) (23). The immunofluorescent images illustrate the successful establishment of a pseudostratified airway epithelium, comprising key cell subtypes of the human airway, signaling that our model had representative cell types at predicted proportions (**Fig 1B-C**).

### Infection of Differentiated Airway Epithelium with *A. fumigatus* and *C. posadasii*

Once validated, our hAECs were infected with *A. fumigatus* or *C. posadasii* to investigate the response of the differentiated airway epithelium to infection. Each ALI culture was incubated with 10^7^ fungal particles of *A. fumigatus* or *C. posadasii* for 6 or 18 hr, respectively. A shorter infection period was required for *A. fumigatus* due to its hyphal formation, which makes the isolation of single cells difficult (24) and the kinetics of cytokine responses seen in hAECs (18). Single cell suspension from the *in vitro* infected hAEC cultures were used to load individual channels on the 10x Chromium Controller and gene expression libraries were subsequently created using the 3’ V3.1 chemistry (25). Pooled libraries were sequenced at a depth of 9,751 or 11,059 reads per cell for *A. fumigatus* vs. mock, respectively. For the *C. posadasii* experiment, the depth sequences were 6,180 or 6,531 reads per cell for infected vs. mock, respectively.

The sequencing data from lung epithelium of both *A. fumigatus* and *C. posadasii* infected samples were analyzed using unsupervised clustering – which allowed us to identify and categorize the relevant cell types – and visualized using uniform manifold approximation and projection (UMAP) (**Fig. 2A-B**). These UMAP plots highlighted the distinct cell populations of the airway epithelium, including basal, club, ciliated, hillock, goblet, and other rarer epithelial subtypes (*e.g.,* ionocytes, tuft, and neuroendocrine cells). The proportion distribution of each cell type was quantified and found to be similar to reported numbers from human airways (26), providing further support for the validity of our model (**Fig. 2C-D**).

**Figure 2.**
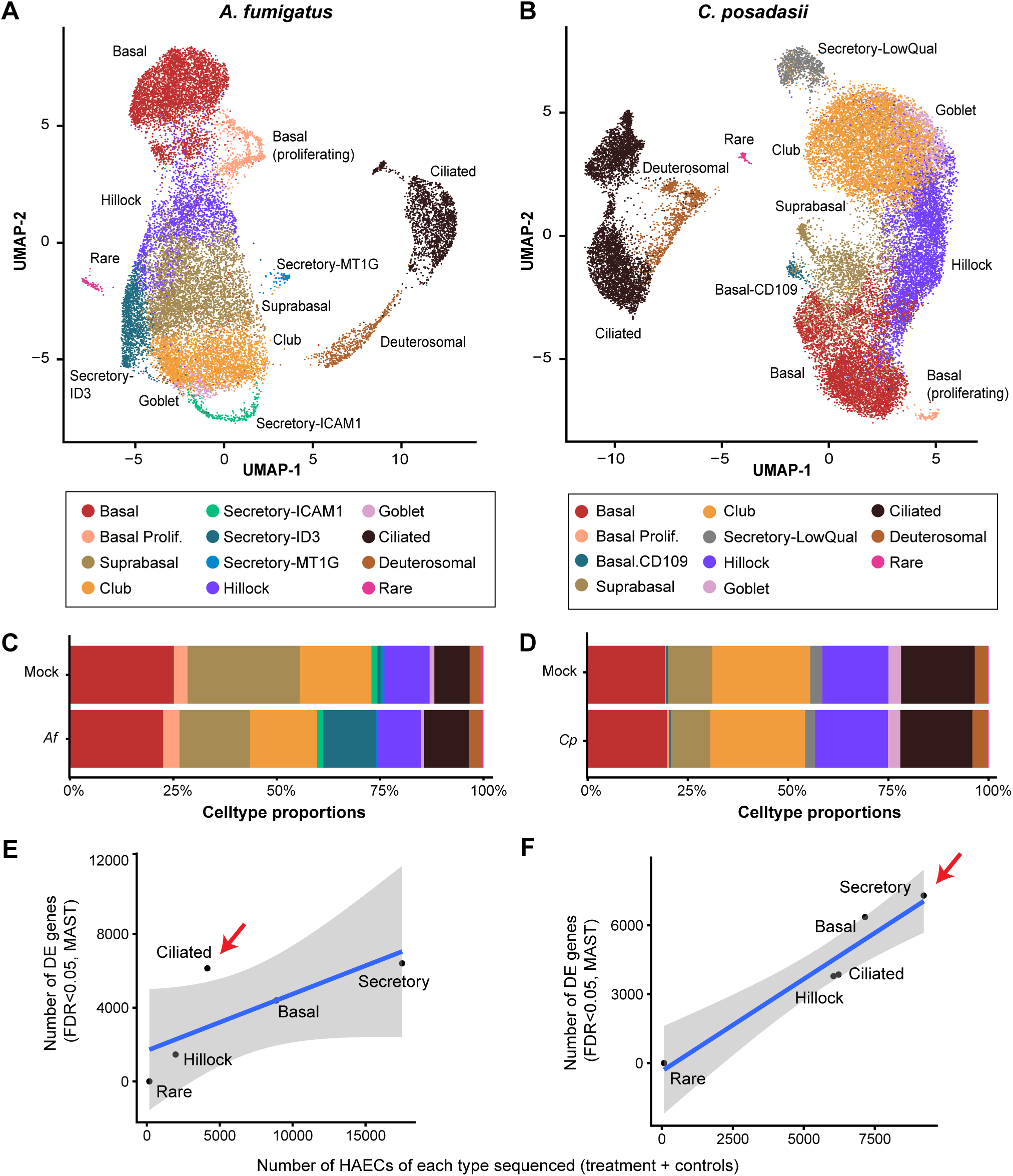
Cell clustering and differential gene expression of scRNA-seq following hAEC infection. UMAP embeddings of 20,560 (hAECs, points) infected with *A. fumigatus* (10^7^ conidia, B5233 strain) **(A)** or 28,724 (hAECs, points) infected with *C. posadasii* (10^7^ arthroconidia, Silveira strain) **(B)** and uninfected controls (mock). Points are colored by assignment to cell types using unsupervised clustering with the Leiden algorithm. Cell type proportions identified during UMAP generation during *A. fumigatus [Af]* **(C)** or *C. posadasii [Cp]* **(D)** infection as compared to mock. Scatter plots showing the relationship between the number of hAECs of each cell type in the scRNA-seq analysis (x-axis) and the number of differentially expressed genes (FDR<0.05) between *A. fumigatus* **(E)** or *C. posadasii* **(F)** and uninfected controls. Blue line: linear model fit, shaded area 90% confidence interval. Red arrows indicate the cell subtype with the highest DEGs. n=1

We then examined the number of differentially expressed genes (DEGs) during infection compared to mock-treated cells. In both infections, we observed little in the way of DEGs in our rare cell subtypes (*i.e.,* ionocytes, neuroendocrine, and tuft cells) because of the low number of hAECs detected in those groups, limiting our statistical power (**Supplemental Fig. 1**). All other cell types, however, displayed a multitude of differentially expressed genes. Our analysis revealed that *A. fumigatus* infection appeared to most dramatically impact ciliated cells, significantly (FDR<0.05) altering the expression of greater than 5,000 genes (**Fig. 2E**), whereas *C. posadasii* infection displayed the greatest effect on secretory cells, inducing significant changes in over 6,000 genes (**Fig. 2F**). This differential response underscores potentially unique cellular targets and mechanisms employed by each pathogen during infection.

### Analysis of Ciliated Cells During *A. fumigatus* Infection

Given that ciliated cells were the most impacted cell type during *A. fumigatus* infection, we performed a detailed analysis of the gene expression profile of these cells compared to mock-treated controls (**Supplemental Figure 2**). Top DEGs were identified using pairwise comparisons with the MAST test. We applied thresholds of a minimum UMI count of 0 and expression in more than 10 cells. The genes were further filtered based on a log_2_ fold change greater than 0.25 and an adjusted p-value (FDR) of less than 0.05. These genes were then visualized using heatmaps, and enrichment analysis was conducted to explore their functional associations. We examined the top-upregulated genes in *A. fumigatus* infected ciliated cells and their corresponding pathways (**Figure 3**).

**Figure 3.**
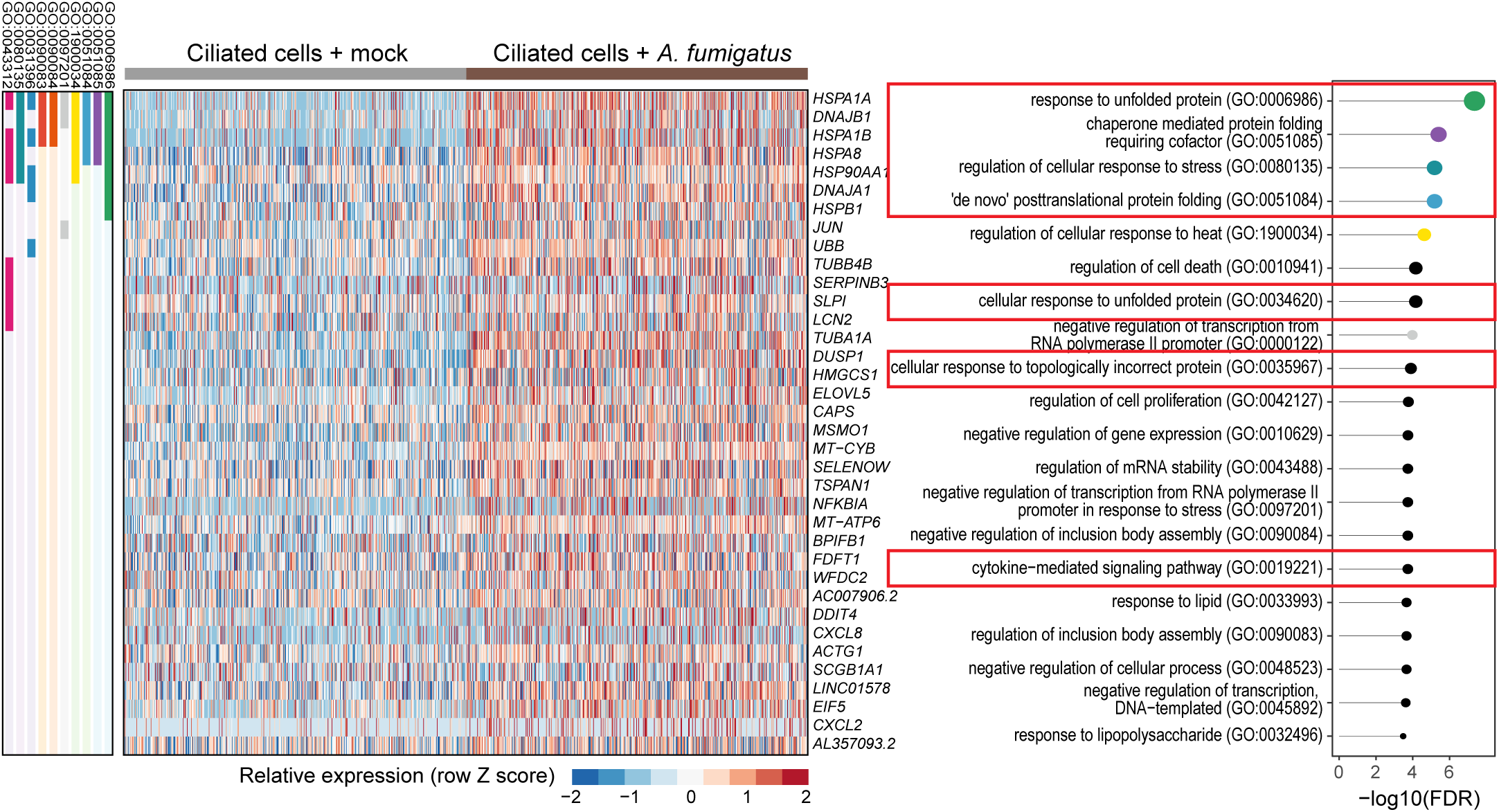
Ciliated cell-specific transcriptional programs activated in hAECs by *A. fumigatus*. Top upregulated (FDR<0.05) genes and gene ontology (GO) biological process pathways in human airway ciliated cells (right columns) by infection with *A. fumigatus*. Individual gene membership in pathways is indicated in the color legend, left. Relevant GO pathways highlighted by the red boxes.

Our analysis revealed a strong enrichment for stress response genes, particularly those associated with the unfolded protein response (UPR). Notably, the genes encoding heat shock proteins such as HSPA1A, HSPA1B, HSPA8, HSP90AA1, DNAJB1, HSPB1, and DNAJA1 were all strongly upregulated. Many of these cytoplasmic chaperones are transcriptionally regulated by the heat shock factor 1 (HSF1) protein, which is sequestered by HSP90 until unfolded proteins compete for HSP90 allowing HSF1 release and non-canonical UPR activation (27). Additionally, the gene *DDIT4*, which is involved in cellular stress response and cell survival, was also highly expressed following stimulation with *A. fumigatus*. This indicates that *A. fumigatus* infection triggers a robust UPR in ciliated cells, likely as a defense mechanism against the pathogen-induced cellular stress.

Moreover, genes encoding key cytokines (*i.e.,* CXCL8, CXCL2) and *NFKBIA* were elevated in infected ciliated cells. CXCL8 and CXCL2 are critical chemokines involved in neutrophil recruitment and activation, which play a vital role in the host’s pro-inflammatory immune response to fungal infections and have previously been shown to be involved in the host response to *A. fumigatus* (17, 18, 28). Innate responses, however, must be tightly regulated to ensure that clearance of the invading pathogen does not lead to unnecessary host cell damage. NFKBIA encodes IκBα, an inhibitor of the NFκB pathway, which regulates inflammation and immune responses as well as cell survival, highlighting potential regulation of this complex immune response (29). Taken together, the increased expression of these cytokines suggests an active inflammatory response to *A. fumigatus* infection, aimed at controlling and clearing the pathogen.

### Analysis of Secretory Cells During *C. posadasii* Infection

We next performed a detailed examination of the transcriptional changes induced by *C. posadasii* infection in secretory cells (**Supplemental Figure 3**) and identified enriched pathways among the top genes upregulated during *C. posadasii* infection (**Figure 4**). Our findings revealed significant transcriptional changes activating the hypoxia response system and immune cell chemotaxis, both of which are hallmarks of *C. posadasii* infection in the lung (30).

**Figure 4.**
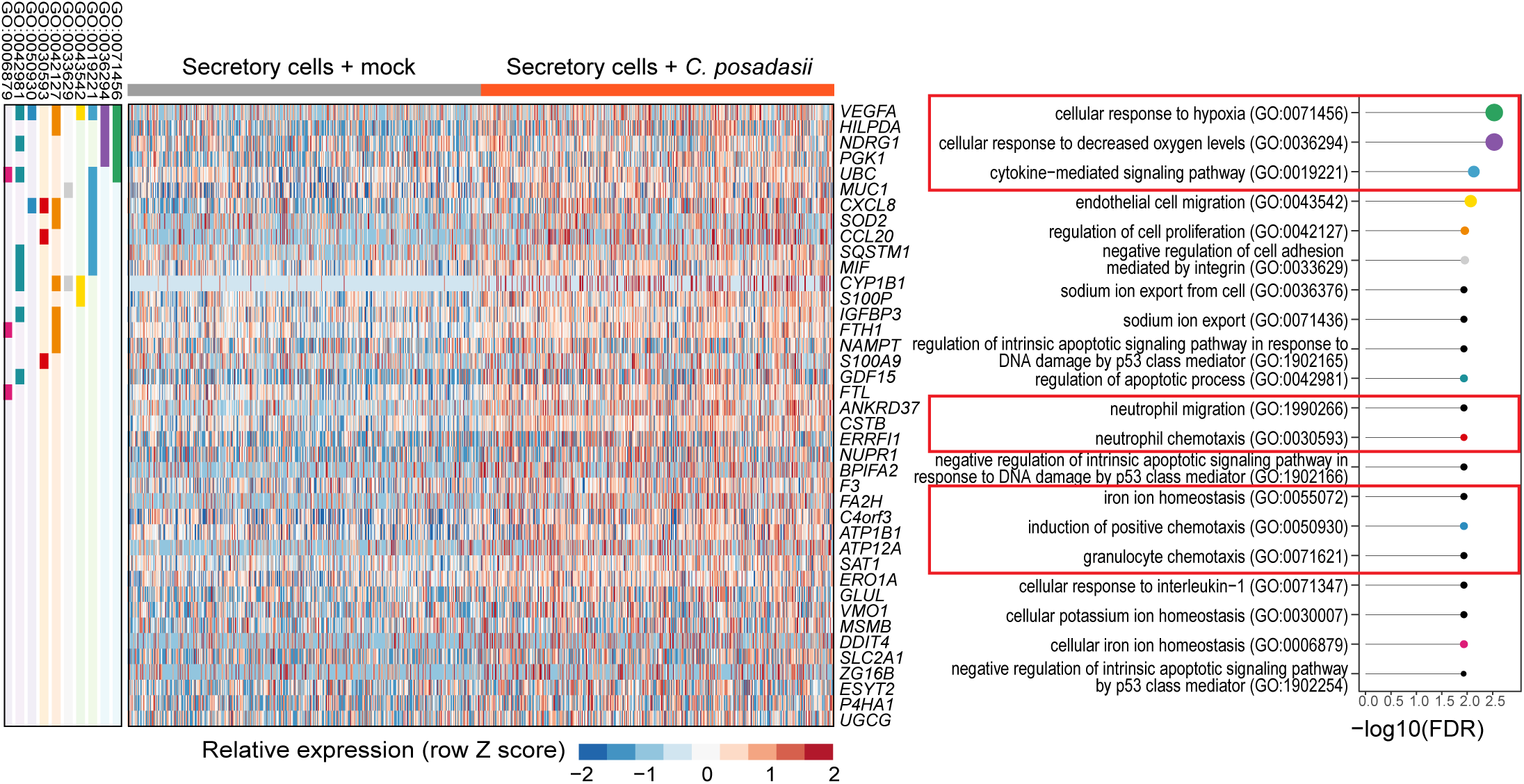
Secretory cell-specific transcriptional programs activated in hAECs by *C. posadasii*. Top upregulated (FDR<0.05) genes and gene ontology (GO) biological process pathways in human airway secretory cells (right columns) by infection with *C. posadasii.* Individual gene membership in pathways is indicated in color legend, left. Relevant GO pathways highlighted by the red boxes.

Many top genes induced by *C. posadasii* are strongly involved in hypoxia and/or cellular stress, highlighting its importance during the early period of infection. VEGFA (vascular endothelial growth factor A) is a critical regulator of angiogenesis and is typically upregulated under hypoxic conditions to promote blood vessel formation and increase oxygen supply (31), a pathway that is very important in the host response to other fungal pathogens (32, 33). Similarly, HILPDA (hypoxia-inducible lipid droplet-associated protein), NDRG1 (N-myc downstream-regulated gene 1), and PGK1 (phosphoglycerate kinase 1) are also all associated with cellular adaptations to low oxygen levels, playing roles in lipid metabolism, stress response, and regulation of anaerobic metabolism, respectively (34–36).

Additionally, the genes FTL (ferritin light chain) and FTH1 (ferritin heavy chain 1) were significantly upregulated, indicating an involvement in ferroptosis and hypoxia. Ferritin is a key regulator of iron homeostasis and plays a role in protecting cells from oxidative stress by sequestering free iron, which can catalyze the formation of reactive oxygen species. The upregulation of FTL and FTH1 suggests a potential mechanism by which *C. posadasii* could be inducing hypoxic stress, namely the disruption of iron homeostasis.

In addition to hypoxia and stress-associated genes, our analyses highlighted the upregulation of several immune-related genes, specifically those involved in regulating immune cell recruitment. Chemokines including CXCL8, CCL20, and MIF recruit immune cells, such as neutrophils, macrophages, and T cells, to sites of infection and inflammation, and play critical roles in the immune response to pathogens (37–40). Furthermore, we identified two members of the S100 family, S100P and S100A9, among the top induced genes in response to *C. posadasii*. The S100 family is a group of calcium-binding proteins that are strongly associated with inflammatory processes and immune cell chemotaxis. Interestingly, CCL20 and other CC chemokines, as well as S100 family members, are heavily induced by hypoxia pathways (41), suggesting a potential link between the specific hypoxic and immune responses induced by *C. posadasii*.

### Common and Differential Gene Expression During *A. fumigatus* and *C. posadasii* Infections

Given the prevalence of immune modulators in both our *A. fumigatus* and *C. posadasii* gene lists, as well as the importance of immune cell recruitment for controlling and clearing fungal respiratory pathogens, we directly compared DEGs encoding paracrine signaling molecules induced by these two pathogens. **Figure 5** illustrates the common and unique differential expression of cytokines and immune regulators across the major cell types (*i.e.,* basal, secretory [goblet & club cells], ciliated, and hillock) of our hAECs during infection with *A. fumigatus* and *C. posadasii*. CXCL8, the most potent human neutrophil attracting chemokine, was the only chemokine robustly induced by both fungi, albeit to differing degrees depending on the cell subtype.

**Figure 5.**
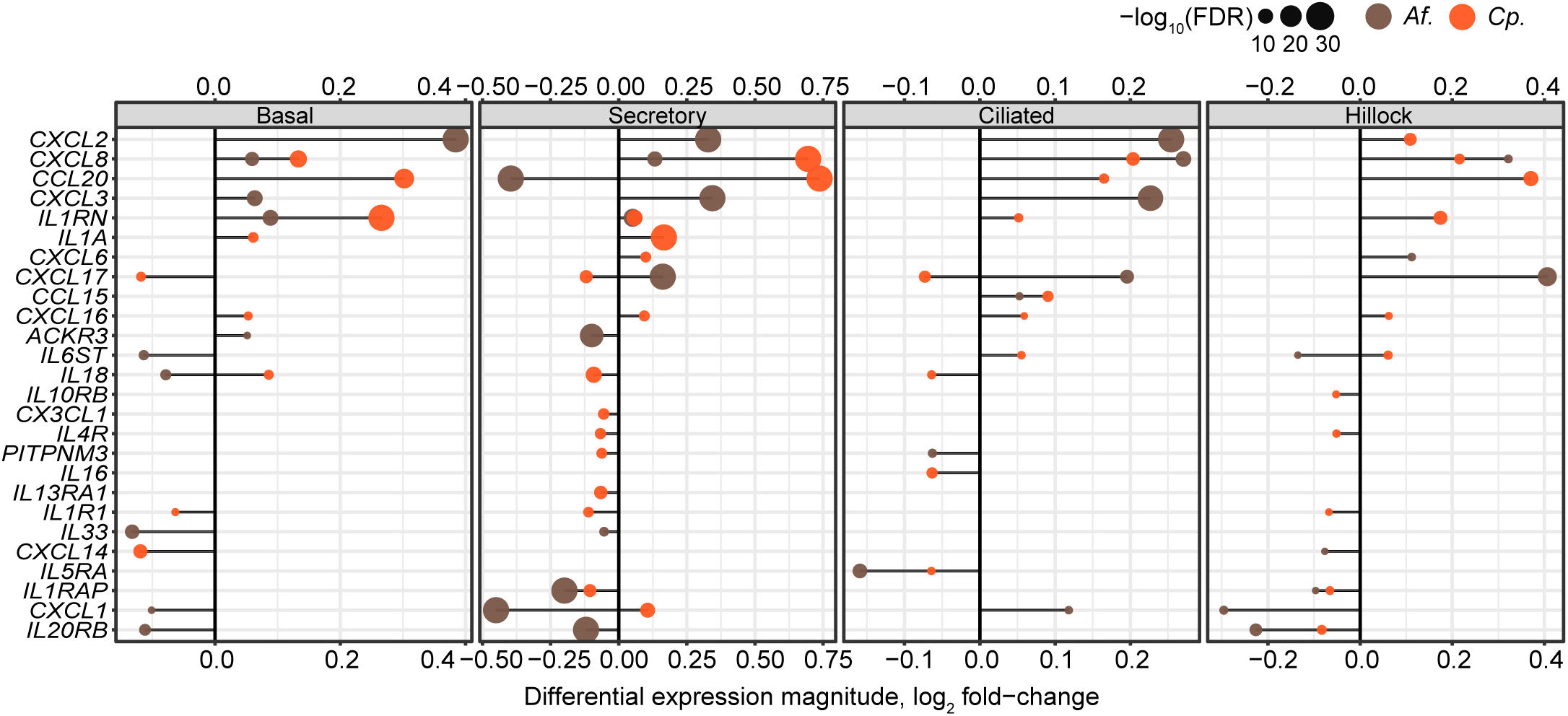
Cell type-specific paracrine signaling responses to fungal pathogens. ‘Lollipop’ plots show the relationship between differential expression effect size (x-axis) and significance (dot size, legend) for any chemokine, interleukin, or cognate receptor (rows, HUGO gene sets189, 483, 601, and 602), by each fungal pathogen (color legend; *Af = A. fumigatus; Cp = C. posadasii*) for each indicated cell type.

Both pathogens displayed robust signatures of cytokine expression involved in the regulation of immune cell recruitment, suggesting that the specific subsets of immune cells recruited might differ between the two fungi. *A. fumigatus* specifically upregulated *CXCL2* and *CXCL3,* in addition to *CXCL8*, in almost all cell types examined. Like CXCL8, both chemokines play significant roles in the recruitment and activation of neutrophils which are essential for the early immune response to fungal infections. Given this unique signature for several key neutrophil-specific chemokines, our data suggest that the lung epithelium is a critical early initiator of the neutrophil response against *A. fumigatus*.

In sharp contrast, *C. posadasii* induced the expression of chemokines that were much less skewed towards neutrophil recruitment. CCL20, also known as MIP-3α, is a far less potent recruiter of neutrophils, instead displaying a stronger preference for lymphocytes and dendritic cells, helping develop adaptive immune responses at the site of infection as opposed to just innate immunity (38, 39, 42, 43). Curiously, we revealed the unique induction of both *IL1α* and *IL1RN* (Interleukin 1 Receptor Antagonist) during *C. posadasii* infection. IL-1α is a potent pro-inflammatory cytokine that not only activates TNFα signaling but also recruits neutrophils to the site of fungal infection (41). IL1RN, however, functions as a direct antagonist to IL-1α, and based on our analysis, the gene is induced at higher levels than *IL1α*. These observations once again point to another piece of evidence where neutrophil recruitment to the lung is uniquely blunted during *C. posadasii* infection as compared to *A. fumigatus*. Overall, these data suggest that infection by *A. fumigatus* or *C. posadasii* does indeed induce unique inflammatory signatures in specific cell subtypes of the lung epithelia that likely impact the outcome and disease presentation.

## Discussion

In this study, we investigated the transcriptional responses of primary hAECs to infections by *A. fumigatus* and *C. posadasii*. By leveraging unbiased, single-cell sequencing approaches, we delineated the cellular and molecular alterations induced by these respiratory fungal pathogens, providing novel insights into the specific pathways and genes involved in the host response. Our results revealed both distinct and shared transcriptional changes elicited by *A. fumigatus* and *C. posadasii* infections. The induction of stress response genes, particularly those associated with the UPR, was a prominent feature of *A. fumigatus* infection. Notably, ciliated cells exhibited the highest number of DEGs, underscoring their vulnerability to fungal assault. *C. posadasii* exhibited a similar, but clearly distinct profile. Secretory cells, not ciliated cells, were most heavily impacted by *C. posadasii*, and the activated cellular stress pathways were strongly associated with hypoxia response as opposed to protein folding. Overall, the initial analysis of the dataset has already identified several interesting avenues of future research that could uncover key insights into the differing mechanisms of controlling and potentially treating these infections.

Perhaps the most obvious difference noted between epithelial immune responses generated to *Aspergillus* compared to *Coccidioides* infection was in chemokine expression. While both pathogens displayed an induction of CXCL8, the strongest chemoattractant for neutrophils, *A. fumigatus* appeared to induce a stronger neutrophil recruitment signature than *C. posadasii* as it uniquely induced CXCL2 and CXCL3. It is well established that both *A. fumigatus* and *C. posadasii* induce neutrophil recruitment to the lung (17, 44–47), but to our knowledge there has not been a direct comparison of the neutrophil recruiting capabilities of the two. It is possible that *A. fumigatus* may be far more potent than *C. posadasii*, and perhaps this discrepancy in neutrophil recruitment explains why healthy patients are able to control *A. fumigatus* infection but not *C. posadasii*.

Another interesting finding was the induction of hypoxia-associated genes by *C. posadasii,* which to our knowledge had not been directly shown before, although one study implicated upregulation of HIF1α in *C. immitis*-resistant mice compared to susceptible mice (30). While we did not detect hypoxic signatures during *A. fumigatus* infection, others have reported it during lung infections at later timepoints than 6 hr (48, 49). The current role in the creation of a hypoxic environment in the lung by fungal pathogens is unclear. Hypoxia regulates both angiogenesis and immune cell signaling/recruitment, vital pathways for controlling dissemination and spread of the pathogen, as well as potentially regulating the fungal lifecycle (33). Furthermore, hypoxic respiratory failure is a rare, but extremely fatal condition that can develop during severe *Coccidioides* infections (50). It is also possible largescale spreading of the pathogen throughout the lung and the induction of this signature contributes to the formation of this severe disorder.

This dataset and our analysis not only enhance our understanding of pulmonary fungal infections but also opens new avenues for research into targeted therapies and interventions and provide opportunities for other investigators to explore pathways within human lung epithelial cells. These data, coupled with recent scRNA-seq datasets from mouse innate immune cells responding to in vivo challenge with *C. posadasii* (51, 52), will enable us to understand the pathogenesis of these invasive fungal diseases. Furthermore, these data provide an exciting new tool for the fungal field to expand their investigation. Future studies will delve deeper into the unexplored aspects of this data, potentially uncovering novel biomarkers and therapeutic targets to improve the management and treatment of fungal respiratory diseases.

## Methods

### Fungal Culture

The *A. fumigatus* B5233 strain was gifted by K.J. Kwon-Chung (National Institutes of Health; NIH) and grown as previously described (18). Briefly, *A. fumigatus* was cultured at 37°C for 3-5 days on glucose minimal medial slants. To harvest conidia, a sterile solution of deionized water with 0.01% Tween 20 was added to each slant, and the surface was gently agitated with a sterile cotton swab. The resulting suspension was filtered through a 40 μm cell strainer to remove hyphal fragments. The conidia were then washed three times with sterile PBS and counted using a LUNA™ automated cell counter. Conidia were used immediately and applied to the apical surface in HBSS media (total volume of 400 µL) (StemCell, #37150).

The *C. posadasii* Silveira strain was received from BEI (NR-48944). WT *C. posadasii* were plated on 2x GYE media (20 g D-(+)-Glucose [Sigma-Aldrich, Cat#G5767], 10 g Bacto Yeast Extract [ThermoFisher, Cat#212750], 15 g Bacteriological agar [Sigma-Aldrich, Cat#A5306], in 1 L of diH_2_O). Cultures were grown at 30°C at ambient CO_2_ for 4-7 weeks with weekly observation. Arthroconidia were harvested using sterile 1x PBS (Corning, Cat# 21-040-CV) and agitated with cell scraper (Corning, Cat#3010). Suspension was passed through 40 µm cell strainer (CellTreat, Cat#229481), vortexed, and centrifuged at 12,000 *g* for 8 mins. The pellet was resuspended in PBS and washed twice. Arthroconidia were used immediately and applied to the apical surface in HBSS media (total volume of 400 µL).

### Isolation and Differentiation of Human Airway Epithelial Cells

Primary hAECs were cultured following established protocols (17, 18). Basal cells were maintained in SAGM (PromoCell, #C-21170), supplemented with 5 μM Y-27632 (Tocris, #1254), 1 μM A-83-01 (Tocris, #2939), 0.2 μM DMH-1 (Selleck Chemicals, #S7146), 0.5 μM CHIR99021 (Tocris, #4423), and 1% penicillin/streptomycin (Gibco, #151410122) on plates coated with laminin-enriched 804G-conditioned media. For differentiation into a pseudostratified epithelium at the ALI, the apical compartments of 12 mm Transwell inserts with 0.4 µm pore polyester membranes (Corning, #3460) were pre-coated with 804G-conditioned media for at least 4 hours. After removing the coating media, a suspension of basal cells in SAGM was applied to the apical compartment, and SAGM was added to the basolateral compartment for overnight incubation. The next day, SAGM was replaced with a 1:1 mixture of PneumaCult-ALI medium (StemCell, #05001) and DMEM/F-12 (Gibco, #11320033) for further overnight incubation. The medium in the apical compartment was then removed to establish the ALI. The hAECs were maintained at ALI for 16-23 days, with the apical compartment kept dry and the basolateral medium refreshed regularly. To ensure intact epithelium, we measured the transepithelial electrical resistance (TEER) on the day of experiment. All epithelium demonstrated a TEER reading of at least 1,000 ohms.

### Infection of hAECs and Isolation of Single Cells

10^7^ infectious units of *A. fumigatus* B5233 or *C. posadasii* Silveira strain were added to the apical side of the hAEC Transwells for 6 or 18 hr, respectively. Once infection was complete, all ALI media was aspirated and hAECs were washed with ice cold wash buffer (1000 µL basolateral, 500 µL apical; 10% heat-inactivated fetal bovine serum [Gibco] in PBS). The wash buffer was pipetted up and down several times to liberate non-adherent particles and repeated twice. After three total washes, the Transwells were transferred to a 50 mL conical containing 6 mL of dissociation solution (TrypLE [Thermo Fisher, #1264013]) and incubated on a rocking platform at 37°C. Every 3-5 minutes conical tubes were vortexed to break up clumps. Once the Transwell had turned clear, dissociation was complete, and the reaction was quenched using one volume of wash buffer. Samples were then passed through a 40 µm filter to remove any large clumps before being spun down at 42 *g* for 5 min. Cells were then resuspended in fresh wash buffer and viability/counts were attained using Trypan Blue (*A. fumigatus* experiment) or acridine orange (*C. posadasii* experiment) and a LUNA™ automated cell counter. Viability was >90%.

### Single-Cell RNA Sequencing and Library Preparation

Single cell suspensions from infected and mock hAECs were loaded into the Chromium Controller (10x Genomics) for droplet generation. For each sample,16,000 cells were loaded per channel aiming for a recovery of 10,000 single cells. There was only one channel per condition. The scRNA-seq libraries were constructed using the Chromium Next GEM single cell 3’ V3.1 Reagent Kit (10x Genomics, PN 1000268). Library quality was assessed with an Agilent 2100 Bioanalyzer and TapeStation. All gene expression libraries were multiplexed and sequenced at the Harvard Biopolymers Core Facility at a depth of 9,751 reads/cell for *A. fumigatus* infected, 11,059 reads/cell for *A. fumigatus* mock, 6,180 reads/cell for *C. posadasii* infected, and 6,531 reads/cell for *C. posadasii* mock on an Illumina Nextseq 500/550 instrument using the high output v2.5 75 cycles kit with the following sequencing parameters: read 1 = 26; read 2 = 56; index 1 = 8; index 2 =0. Demultiplexing the sequence reads to create FASTQ files and alignment to the human genome reference GRCh38 (version refdata-gex-GRCh38-2020-A, 10X Genomics) were performed using Cell Ranger (version 7.1.0, 10X Genomics) commands mkfastq and count, respectively, and subsequent analysis was performed to evaluate transcriptional changes.

### Data Analysis

Quality filtering, variable gene selection, and clustering were performed as described previously (53). We interpreted clusters using known markers of airway epithelial subtypes, merging clusters expressing the same markers. For each cell, we quantified the number of genes for which at least one read was mapped, and then excluded all cells with fewer than 800 or greater than 10,000. We also excluded cells in which more than 30% of transcripts mapped to the mitochondrial genome. Expression values *E_i_*_,*j*_ for gene *i* in cell *j* were calculated by dividing UMI count values for gene *i* by the sum of the UMI counts in cell *j*, to normalize for differences in coverage, and then multiplying by 10,000 to create TPM-like values, and finally calculating log_2_(TPM+1) values.

For each gene, we modeled the relationship between detection fraction (proportion of cells in which at least one UMI was observed) and the log of total number of UMIs using logistic regression. Outliers from this curve are expressed in a lower fraction of cells than would be expected, and are thus highly variable, that is, they are specific to a cell-type, treatment, condition, or state. We selected the top 3,000 genes with the highest residuals as highly variable genes. We restricted the expression matrix to the subsets of variable genes and high-quality cells noted above, and values were centered and scaled before input to PCA. For *C. posadasii*, data from different conditions were then integrated using the Harmony algorithm (54), before shared nearest-neighbor network construction and clustering using the Leiden algorithm as we have described previously (55). Differential expression (DE) tests were performed using MAST (56). All DE tests were run by comparing all cells of each type between conditions. For each cell type, genes were only tested if they were detected in greater than 10 cells. Chemokines were selected using the HUGO gene set chemokine ligands (group 483, https://www.genenames.org/data/genegroup/#!/group/483). Pathway enrichment was performed using the ‘EnrichR’ R package.

## Data Availability Statement

The raw data associated with scRNA-seq studies are available in the GEO database at [ascension number not assigned yet].

## Acknowledgments

The following reagent was obtained through BEI Resources, NIAID, NIH: *Coccidioides posadasii*, Strain Silveira, NR-48944. This work was supported by NIH grant 1K08AI14755 (J.L.R); R01AI168370, R01AI040996, R37AI035681 (B.S.K); R01HL164563 (J.R.); K08HL157725 (P.S.); R01AI150181, R01AI136529, R21AI152499 (J.M.V); and an American Heart Association Career Development Award (P.S.)

## Abbreviations

ALI: air-liquid interface
DEGs: differentially expressed genes
DE: differential expression
FTH1: ferritin heavy chain 1
FTL: ferritin light chain
hAECs: human airway epithelial cells
HSF1: heat shock factor 1
IL1RN: interleukin 1 receptor antagonist
NDRG1: N-myc downstream-regulated gene 1
PGK1: phosphoglycerate kinase 1
SAGM: small-airway epithelial cell medium
scRNA-seq: single-cell RNA sequencing
UMAP: uniform manifold approximation and projection
UPR: unfolded protein response
VEGFA: vascular endothelial growth factor A

